## Supplementary figures and images for "Repression of PRMT activities sensitize homologous recombination-proficient ovarian and breast cancer cells to PARP inhibitor treatment"

### Figure S1

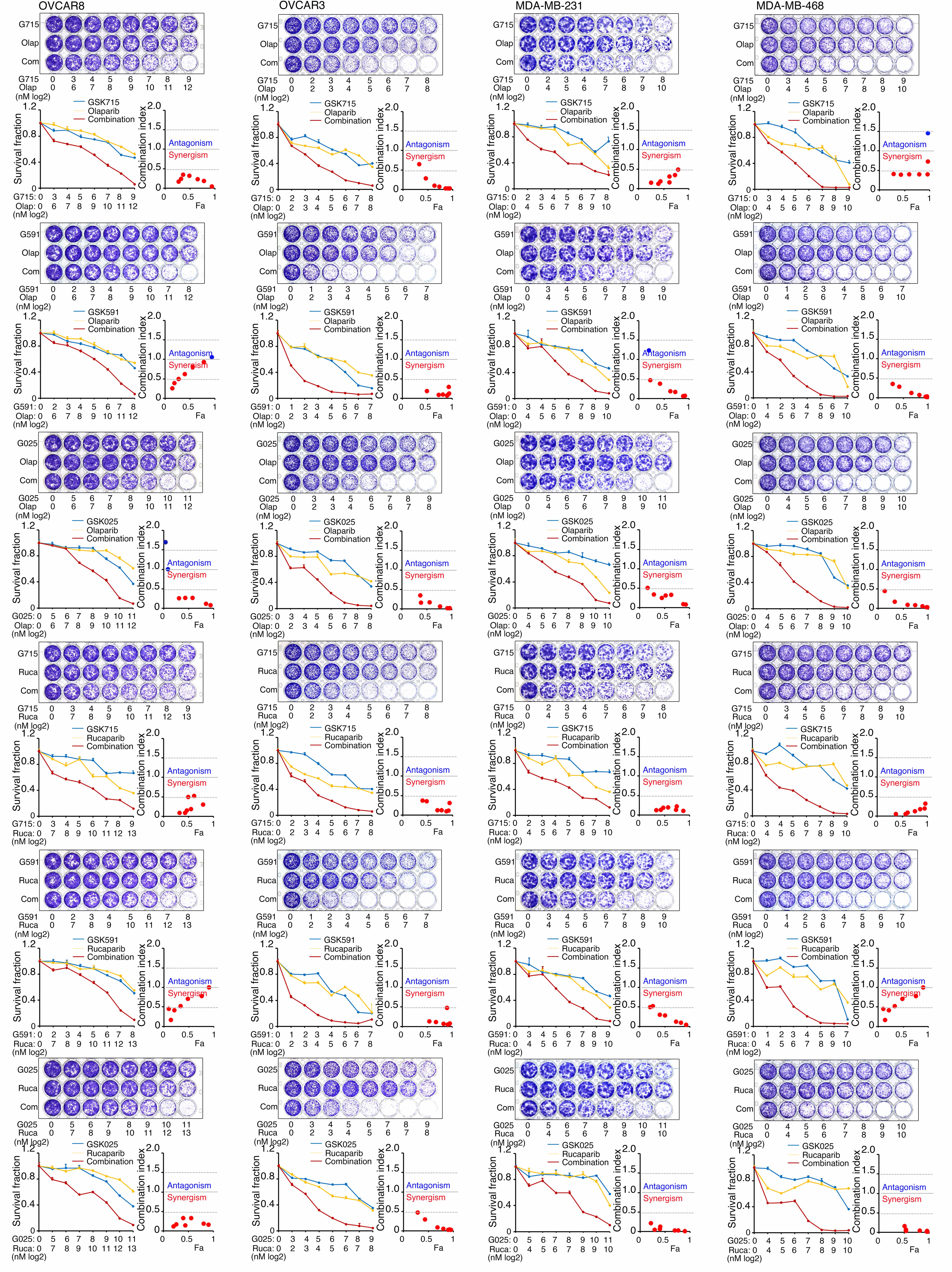

### Figure S2

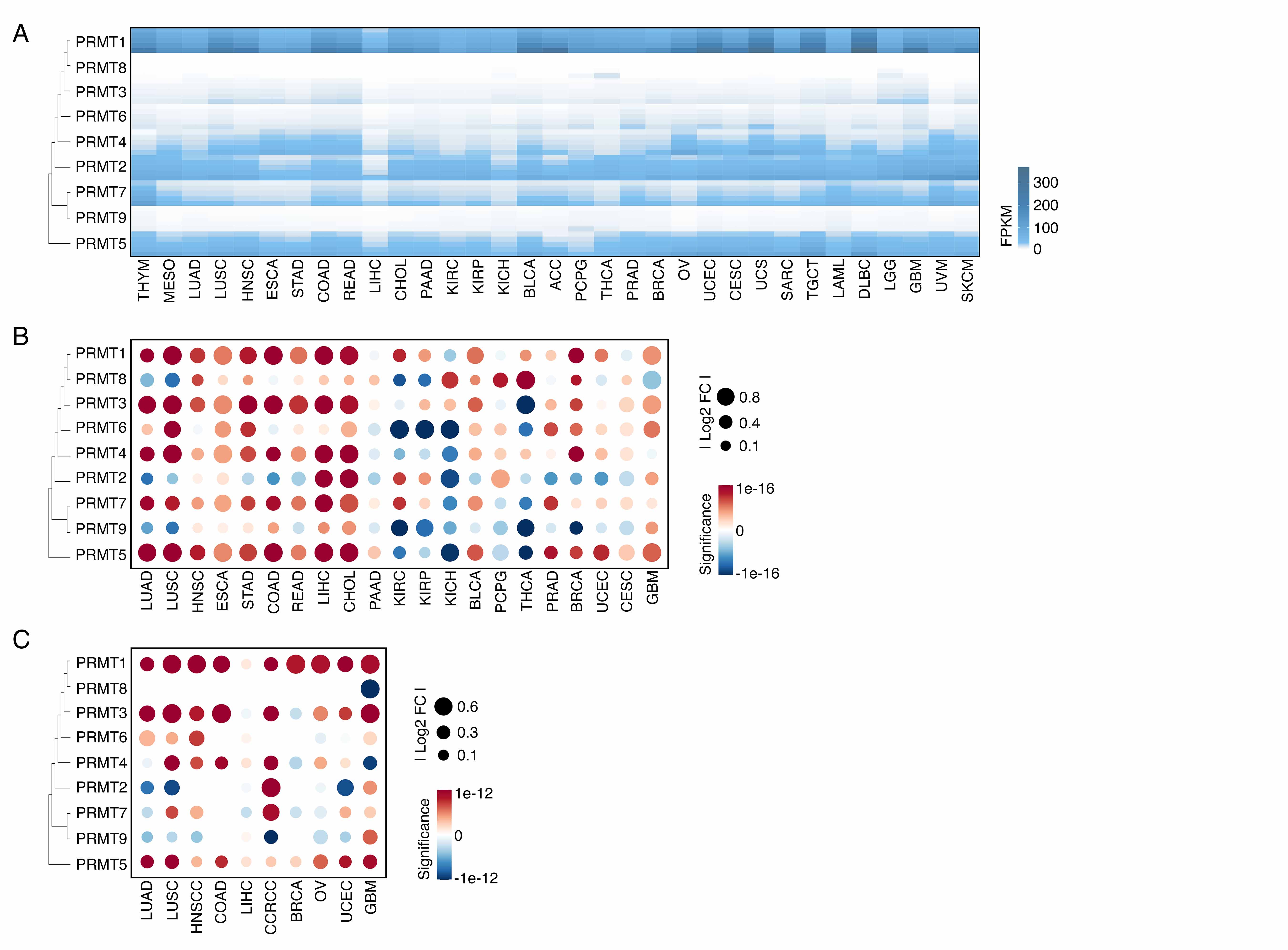

### Figure S3

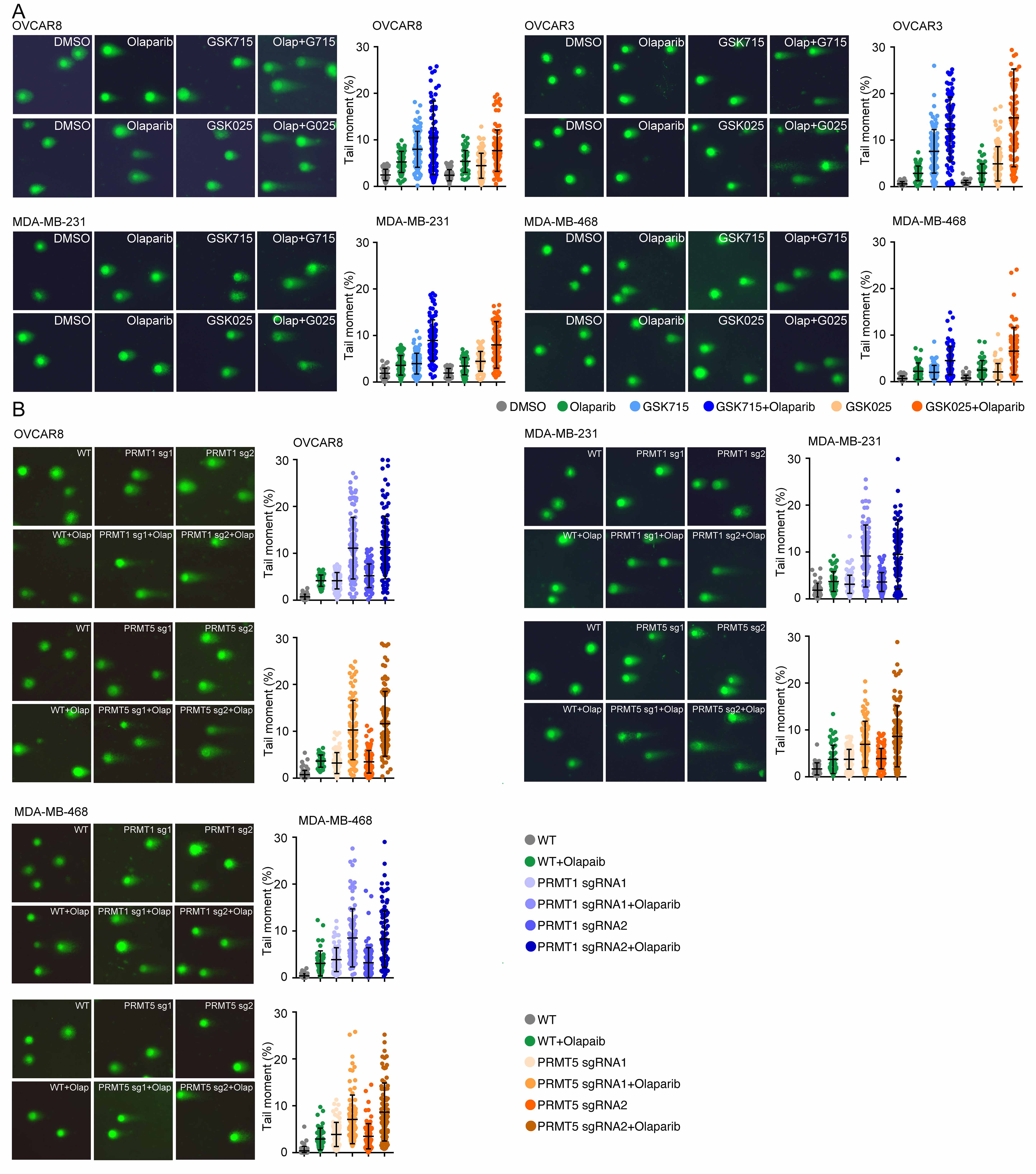

### Figure S4

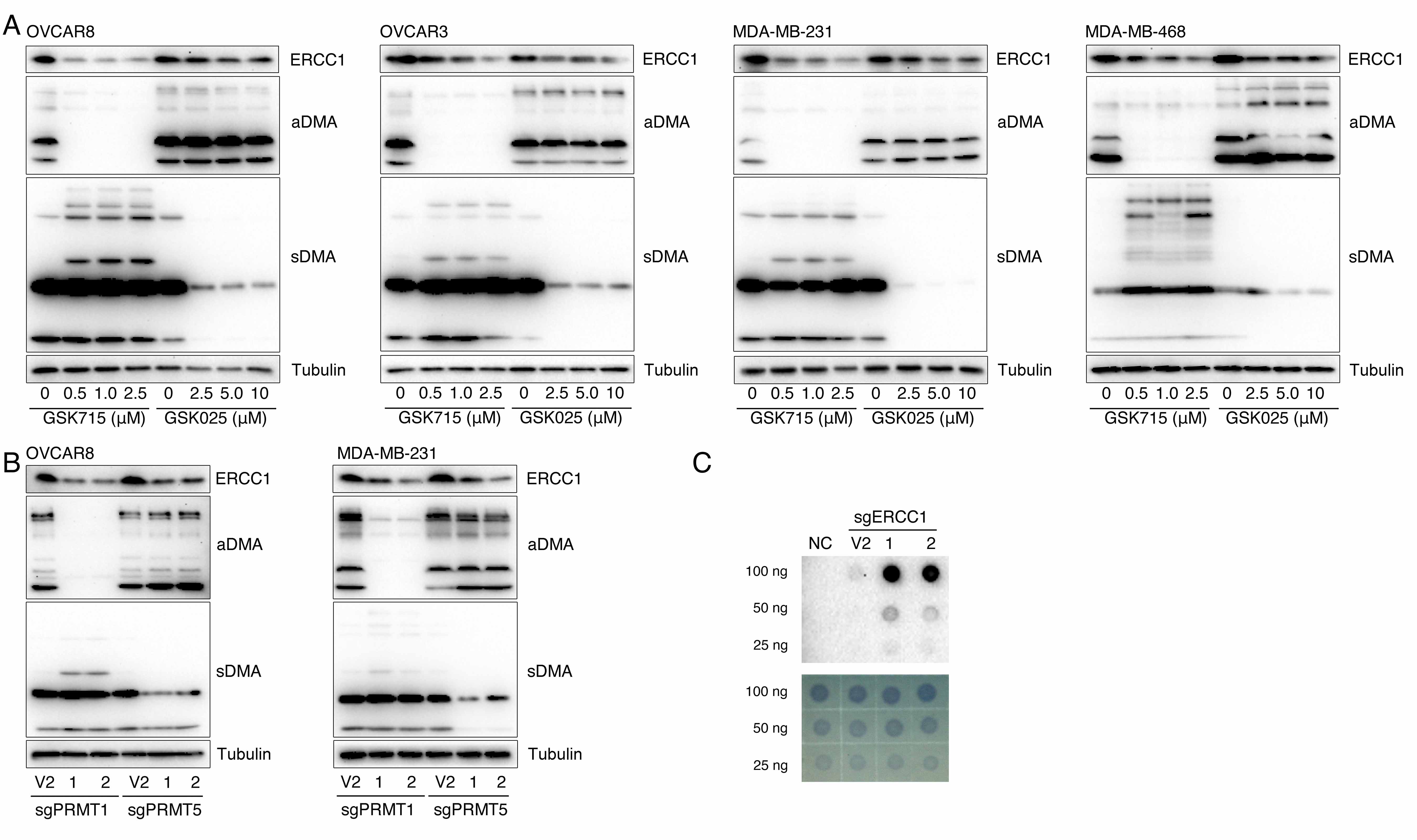

### Figure S5

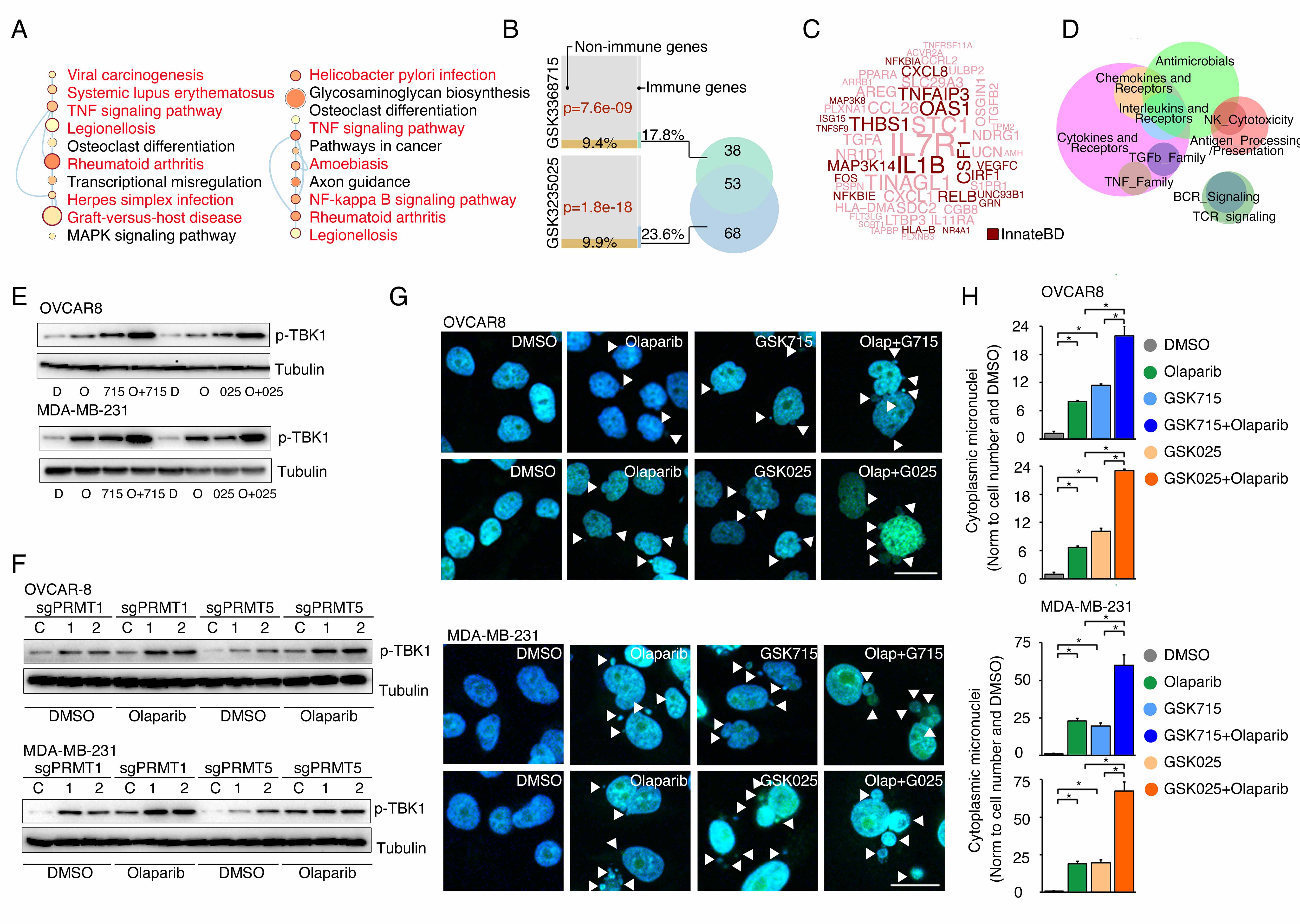

### Figure S6

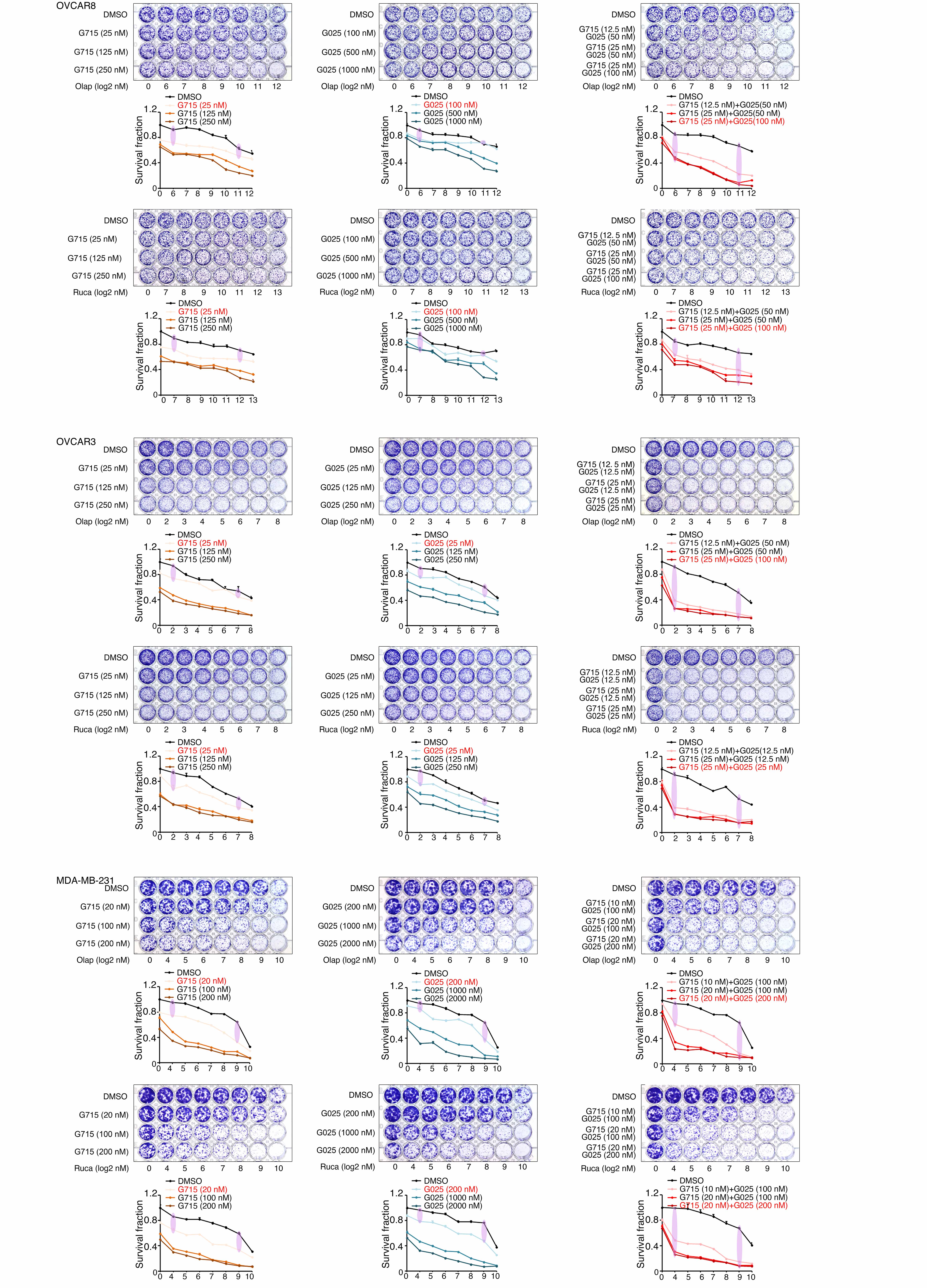
